# Nonionic surfactants can modify the thermal stability of globular and membrane proteins interfering with the thermal proteome profiling principles to identify protein targets

**DOI:** 10.1101/2022.10.17.512546

**Authors:** Emmanuel Berlin, Veronica Lizano-Fallas, Ana Carrasco del Amor, Olatz Fresnedo, Susana Cristobal

## Abstract

The membrane proteins are essential targets to understand cellular function. The unbiased identification of membrane protein targets is still the bottleneck for a system-level understanding of cellular response to stimuli or perturbations. It has been suggested to enrich the soluble proteome with membrane proteins by introducing nonionic surfactants in the solubilization solution. This strategy was aiming to simultaneous identify the globular and membrane protein targets by thermal proteome profiling principles. However, the thermal shift assay would surpass the cloud point temperature from the nonionic surfactants frequently utilized for membrane protein solubilization. It is expected that around the cloud point temperature, the surfactant micelles would suffer structural modifications altering protein solubility. Here, we show that the presence of nonionic surfactants can alter protein thermal stability from a mixed, globular and membrane, proteome. In the presence of surfactant micelles, the changes in proteins solubility analyzed after the thermal shift assay were affected by the thermal dependent modification of the micellar size, and its interaction with proteins. We demonstrate that the introduction of nonionic surfactants for the solubilization of membrane proteins is not compatible with the principles of target identification by thermal proteome profiling methodologies. Our results lead to explore thermal-independent strategies for membrane protein solubilization to assure confident membrane protein target identification. The proteome-wide thermal shift methods have already shown their capability to elucidate mechanisms of action from pharma, biomedicine, analytical chemistry, or toxicology and finding strategies, free from surfactants, to identify membrane protein targets would be the next challenge.

## INTRODUCTION

Understanding the consequence of the interaction of membrane proteins with small molecules such as drugs or chemicals would require from high-throughput methodologies supporting a rapid screening of several thousand protein candidates from any cell type. The membrane proteins are essential targets to understand cellular function. They are key nodes that regulate the cell communication and initiation of signal transduction pathways. Membrane proteins are largely represented among the drug targets for its capability to orchestrate the cellular responses^1^. Moreover, the membrane proteins are primary interactors with small chemicals present in extracellular compartments, could facilitate the cellular uptake^2^, and could be responsible of the molecular initiation events leading to adverse outcome pathways for human health^3^.

Different methodologies have been applied for membrane protein target identification. In general, there were methods with low throughput capability, and a sufficient similarity to previously described targets as a prerequisite for new findings^1^. Important changes in the field of target engagement arise with the introduction of the protein thermal shift assay in a proteome-wide context. The Cellular thermal shift assay relayed on a large collection of antibodies for protein identification, and quantitation^4^. However, the introduction of mass spectrometry (MS) to analyze the thermal-induced changes of proteome solubility was pivotal to enable the unbiased identification of targets from soluble proteome in methods called Thermal proteome profiling (TPP)^5^, and later Proteome integral solubility alteration^5,6^. The developments of these methodological approaches have recently explored many scientific problems defined by interactions of small chemicals and cellular proteins from biomedicine, analytical chemistry^7^. At our lab, we have focused on implementation for its application to biodiscovery^8^ or toxicology^9^.

Searching for targets in proteomes containing globular proteins facilitates the application of this methodology because the protein-chemical interaction is the main or only stimulus that induces perturbation in protein solubility under the thermal treatment. The factors affecting and ruling protein solubility enlarge when the studied proteome contains a mixture of globular and membrane proteins. Therefore, the next challenge was how to implement those methods for the identification of membrane protein targets. The reported strategy has been aiming to enlarge the soluble proteome with the addition of nonionic surfactants to the solubilization buffer. The surfactants tested are routinary used in membrane protein studies, such as nonyl-phenyl-polyethylene glycol (NP-40S)^10^, or Igepal CA-630 (Igepal), composed of octyl-phenoxy(polyoxyethylene)ethanol ^11^.

Those nonionic surfactants are substitutes for the original octyl-phenoxy(polyoxyethylene)ethanol (NP-40) that although has been widely used, it is no longer produced. The chemical structure of Igepal resembles better the original NP-40 formulation than the new NP-40S. However, NP-40S offers closer values to NP-40 in parameters that are important for the micelle formation, such as hydrophile-lipophile balance or molecular weight^12^. The cloud point temperature (CPT) is one of the most distinct physical properties of nonionic surfactants. It is the temperature above which a micellar solution of nonionic surfactant forms spontaneously a two-phase separation. The aqueous micellar two-phase system has been frequently used for fractionation of proteins with different hydrophobicity^13^. For NP-40 and NP-40S, the CPT occurs at 45-50 °C and at 53-67 °C for Igepal. Looking into the TPP workflow for identification of membrane proteins, it is important to observe that the thermal shift spans from 37-67 °C. This temperature range includes the specific CPTs of the nonionic surfactants used to increase the membrane protein solubilization ^10^.

The expected temperature-dependent modifications of the solution with micelles of nonionic surfactants would include at least micellar stratification^14^. Additionally, during the increased temperature and achievement of CPT, the surfactant micelles are dehydrated acquiring an enlarged and elongated shape^15^. Over the CTP, it was reported that micelle-embedded proteins will be oversaturated in the micelle-rich layer while non-micelle bound proteins exist in the aqueous-rich layer^16^. This may cause crowding, protein-protein interactions, micelle-protein interactions, micelle aggregation, diverse changes in the local environment of the proteome which may affect protein stability^17^.

Given the numerous studies based on the solubilization of membrane proteins with nonionic surfactant, it has been assumed that the membrane proteome in micelles of nonionic surfactants could be compatible with the TPP principles^10^. The TPP principles for the identification of protein targets is solely based on unique alteration on protein solubility, specifically, the beneficial effects of protein-chemical interactions to increase protein thermal tolerance^5^. Therefore, insufficient attention has been paid to evaluate the modification of the aqueous micellar solution at temperature close or over surfactant CPT, and if the structural and density changes on the solvent solution could contribute to alter the protein stability in solution independently of any protein-chemical interaction. This evaluation would offer new insights to define if the chemical-protein interaction in solubility is still a robust parameter to identify membrane protein targets in a solution containing nonionic surfactant micelles that are very malleable within the thermal shift range of temperatures.

In this study, we evaluate the dynamics of nonionic surfactant micelles along with the thermal shift methodology and evaluate temperature dependent alteration of the solubility of the soluble and membrane proteome. The interactions of nonionic surfactants with the soluble and membrane proteome could alter their thermal stability interfering with the proteome-wide thermal shift principles. Considering the urgency to offer high-through methodologies for the unbiased identification of membrane protein targets, this study aims to provide the factors and constraints for the future implementation of TPP for identification of membrane proteins.

## MATERIALS AND METHODS

### Collection of liver tissue

Liver from a 2-month-old female, Sprague Dawley rats were obtained from the University of the Basque Country with an ethics approval code of M20/2016/237. The procedures conducted on the animal were approved by the Ethics Committee for Animal Welfare of the University of the Basque Country UPV/EHU and followed the EU Directives for animal experimentation.

### Nonionic surfactants in the study

The nonionic surfactants and properties utilized in this study were octyl-phenoxy(polyoxyethylene)ethanol, named as Igepal CA-630 (Igepal), and nonyl-phenyl-polyethylene glycol, named as Nonidet-P40 substitute (NP-40S) (Table 1).

**Table 1.** Overview of nonionic surfactant properties: Igepal CA-630 (Igepal), and Nonidet-P40 substitute (NP-40S).

| Surfactant | Chemical formula | HLB | Mw (g/mol) | CPT (°C) |
| --- | --- | --- | --- | --- |
| Igepal | octyl-phenoxy(polyoxyethylene)ethanol | 13 | 603 | 53-67 |
| NP-40S | nonyl-phenyl-polyethylene glycol | 13.5 | 680 | 45-50 |

HLB: hydrophilic lipophilic balance, Mw: molecular weight

### Thermal proteome profiling (TPP). Protein extraction

Liver tissue was resuspended in buffer containing PBS (control), 0.4 % (v/v) Igepal or 0.4 % (v/v) NP-40S in PBS. Tissue samples were mechanically homogenized using a TissueLyser (Qiagen) during 3 min at 25 Hz. Previously, zirconium oxide beads were added to each sample at a ratio 1:1 (v/v). Subsequently, the unbroken cells were lysed by sonication in cycles of 10s/5s for 3 min at 6–10 μm amplitude at 50% intensity from an exponential ultrasonic horn of 3 mm in a Soniprep 150 MSE (MSE Ltd., Lower Sydenham, London, UK). The insoluble fraction was sedimented by centrifugation at 100,000g for 60 min at 4 °C^8^. The soluble proteome was used to perform the Thermal proteome profiling assay. The pellets were lysed with 100 μl of RIPA buffer (RIPA Lysis Buffer, BOSTER) to perform protein identification and quantification of the insoluble fraction. Protein concentration was determined by BCA assay^18^.

### Thermal shift assay (TSA)

The soluble fractions of the three conditions (control, Igepal, and NP-40S) were heated by duplicate at the specific ten temperatures selected for the thermal shift assay: 37 °C, 42 °C, 46 °C, 49 °C, 51 °C, 53 °C, 55 °C, 58 °C, 62 °C, and 67 °C. Aliquots containing 50 μg of protein were independently heated at the corresponding temperature for 3 min, followed by 3 min at room temperature using a thermocycler (MJ Mini Personal Thermal Cycler PTC 1148, Bio-Rad) Samples were centrifugated at 100 000 g for 20 min at 4 °C.

### Ultrasonic-Based Filter Aided Sample Preparation

All the samples analyzed by MS were digested following this method ^**19**^. Briefly, the proteins were reduced by adding 200 μL of 50 mM dithiothreitol (DTT) prepared in 8 M urea and 25 mM ammonium bicarbonate (AmBic) in 30 kDa microcon centrifugal filter units. Ultrasound energy was then applied using the ultrasonicator Q700 Sonicator with cup horn (Qsonica L.L.C) for 5.25 min (7 cycles: 30 s on and 15 s off UT, 25% UA, 20 kHz UF). Afterward, centrifugation for 20 min at 14 000 g was done, followed by protein alkylation by the addition of 100 μL of 50 mM iodoacetamide (IAA) prepared in 8 M urea and 25 mM AmBic solution. The alkylation step was sped up using the ultrasonicator Q700 Sonicator with cup horn during 5.25 min (7 cycles: 30 s on, and 15 s off UT, 25% UA, 20 kHz UF). Finally, 100 μL of 1:30 trypsin in 12.5 mM AmBic solution was added, and the protein digestion was processed using the ultrasonic microplate horn assembly for 5.25 min (7 cycles: 30 s on, and 15 s off UT, 25% UA, 20 kHz UF). To ensure that all the peptides were extracted, 100 μL of 3% (v/v) acetonitrile (ACN) and 0.1% (v/v) formic acid (FA) were added followed by a centrifugation of 20 min at 14 000 g. This previous step was repeated one more time. The samples were acidified with 10% FA to achieve pH between 3 and 2. The desalting process was performed by reverse phase chromatography in C18 top tips using ACN (60% v/v) with FA (0.1% v/v) for elution, and vacuum dried to be stored at −80 °C till further analysis.

### Nano LC-MS/MS analysis

The desalted peptides were reconstituted with 0.1% FA in ultra-pure milli-Q water and the concentration was measured using a Nanodrop (Thermo Scientific). Peptides were analyzed in a QExactive quadrupole orbitrap mass spectrometer (Thermo Scientific). Samples were separated using an EASY nLC 1200 system (Thermo Scientific) and tryptic peptides were injected into a pre-column (Acclaim PepMap 100 Å, 75 um × 2 cm) and peptide separation was performed using an EASY-Spray C18 reversed-phase nano LC column (PepMap RSLC C18, 2 um, 100 Å, 75 um × 25 cm). A linear gradient of 6 to 28% buffer B (0.1% FA in ACN) against buffer A (0.1% FA in water) during 78 min, followed by 40% buffer B against buffer A till 95 min was applied. The last 5 min (95-100 min) had a cleanse period with 100% B solution. The linear gradient was carried out with a constant flow rate of 300 nL/min. Full scan MS spectra were recorded in the positive mode electrospray ionization with an ion spray voltage power frequency (pf) of 1.9 kV (kV), a radio frequency lens voltage of 60 and a capillary temperature of 275 °C, at a resolution of 30,000 and top 15 intense ions were selected for MS/MS under an isolation width of 1.2 m/z units. The MS/MS scans with higher energy collision dissociation fragmentation at normalized collision energy of 27 % to fragment the ions in the collision induced dissociation mode.

### Peptide and protein identification and quantification

The proteins were identified using Proteome Discoverer (version 2.1, Thermo Fisher Scientific) The MS/MS spectra (. raw files) were searched by Sequest HT against the *Rattus norvegicus* database from Uniprot (UP000002494; 47,954 entries). A maximum of 2 tryptic cleavages were allowed, the precursor and fragment mass tolerance were 10 ppm and 0.6 Da, respectively. Peptides with a false discovery rate (FDR) of less than 0.01 and validation based on q-value were used as identified. The minimum peptide length considered was 6 and the FDR was set to 0.1. Proteins were quantified using the average of top three peptide MS1-areas, yielding raw protein abundances. Common contaminants like human keratin and bovine trypsin were also included in the database during the searches for minimizing false identifications. The mass spectrometry proteomics data have been deposited to the ProteomeXchange Consortium via the PRIDE partner repository with the dataset identifier PXD037153^20^.

### Protein localization

The bioinformatic tool PANTHER^21^, which utilizes Gene Ontology classification, was used for protein localization. To note, one protein may have several sub-localizations. This selection was made by choosing proteins tagged with the GO term: GO:0016020.

### Thermal proteome profiling analysis

Melting curves were calculated using a sigmoidal fitting approach with the R package TPP ^22^. The melting curves were fitted after normalization following the equation described ^5,22^ computed in R:

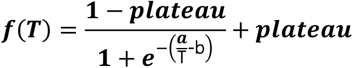

where T is the temperature, and a, b and “plateau” are constants. The value of f(T) at the lowest temperature Tmin was fixed at 1. The melting point of a protein is defined as the temperature Tm at which half of the amount of the protein has been denatured. The quality criteria for filtering the sigmoidal melting curves were: (i) fitted curves for both vehicle- and compound-treated conditions had an R^**2**^ of >0.8; (ii) the vehicle curve had a plateau of <0.3; (iii) the melting point differences under both the control and the treatment conditions were greater than the melting point difference between the two controls; and (iv) in each biological replicate, the steepest slope of the protein melting curve in the paired set of vehicle- and compound-treated conditions was below −0.06. The NPARC of the R package was used to detect significant changes in the temperature-dependent melting behavior of each protein due to changes in experimental conditions^22^. The significance threshold was set at p < 0.05.

### Dynamic light scattering (DLS)

DLS measurement were obtained from the surfactant solutions studied here after exposed to 10 stepwise increases of temperatures. Solutions were PBS as control, 0.4 % Igepal and 0.4 % NP-40S. The temperatures chosen were those assayed in the TSA: 37 °C, 42 °C, 46 °C, 49 °C, 51 °C, 53 °C, 55 °C, 58 °C, 62 °C, and 67 °C. A 384-well plate was used and for each temperature a volume of 40 μL of each sample was put into the wells. None of the samples contained proteins due to interference with the DLS reading. To reduce condensation, a silicon solution was added on top of each well. The DLS (DynPro II Platereader, Wyatt) was set to increase the plate temperature following the TSA protocol and for each temperature, 10 acquisitions time (sec) were performed. Images were taken between the temperatures 37 °C to 49 °C since temperatures above were too high for the camera to operate safely. During this thermal exposure both readings of micelle size, and images were taken. Another DLS experiment was performed to measure changes in the DLS measurements from samples heated at 55 °C and 67 °C (CPTs) and then cooled to 21 °C, similarly as the TPP method that includes that cooling step at R.T. for 3 min before ultracentrifugation. All experiments were performed in triplicates.

### Microscopy

Visualization of samples exposed to heat were performed to detect surfactant and/or protein aggregates by preparing the samples following the TPP method. Control, Igepal, and NP-40S conditions in a volume of 210 μL each were subjected to 67 °C for 3 min in a thermocycler. This temperature was set to ensure that both, Igepal and NP-40S, samples have passed their cloud point. After 3 min of heating, 200 μL of each sample was put into a glass bottom plate (P35-G-1.5-10-C, MatTek) and a cover glass was laid on top of the well. A Leica DMi9 microscope was used with a mounted heating box set at 51 °C (maximum temperature of the heating box) to slow down the cooling of the sample. Visualization was performed within the first 3 min using a HC PL APO CS2 63x/1.20 WATER UV objective with a TL-DIC contrast for 10 ms exposure for each image.

## RESULTS AND DISCUSSION

### Solubilization of membrane proteins for TPP analysis

The introduction of nonionic surfactants in the extraction solution of proteins aims to increase identification of membrane protein targets by TPP without extending the steps and complexity of the extraction procedures. We evaluated differences in the composition and distribution of protein classes in the soluble proteome extracted under native conditions with aqueous solution of PBS containing nonionic surfactant Igepal or NP-40S in comparison to the PBS extracted proteome as control. Those extracting solutions aim to maintain the proteins in native state, a requirement from TPP methodologies. Here we analyzed by quantitative proteomics the soluble proteome, sample that could further utilize for TPP and the insoluble fraction collected in the pellet after centrifugation. The soluble fractions were analyzed and classified according to subcellular localization categories of GO. The number of proteins in each hit was normalized against the total number of proteins. The proteins analyzed were retrieved by choosing proteins present in two out of three replicates. The number of membrane proteins solubilized in the different solutions were similar, 23% in PBS (1550 proteins), 24% in Igepal (1618 proteins), and 22% in NP-40S (1403 proteins). However, looking at specific GO categories that contain integral membrane proteins such as leaflet of the membrane bilayer, and side of membrane component of the membrane, NP-40S increased 50%, and Igepal 20% in the number of these membrane proteins in the soluble fraction compared to control (Fig. 1. A, B).

**Figure 1.**
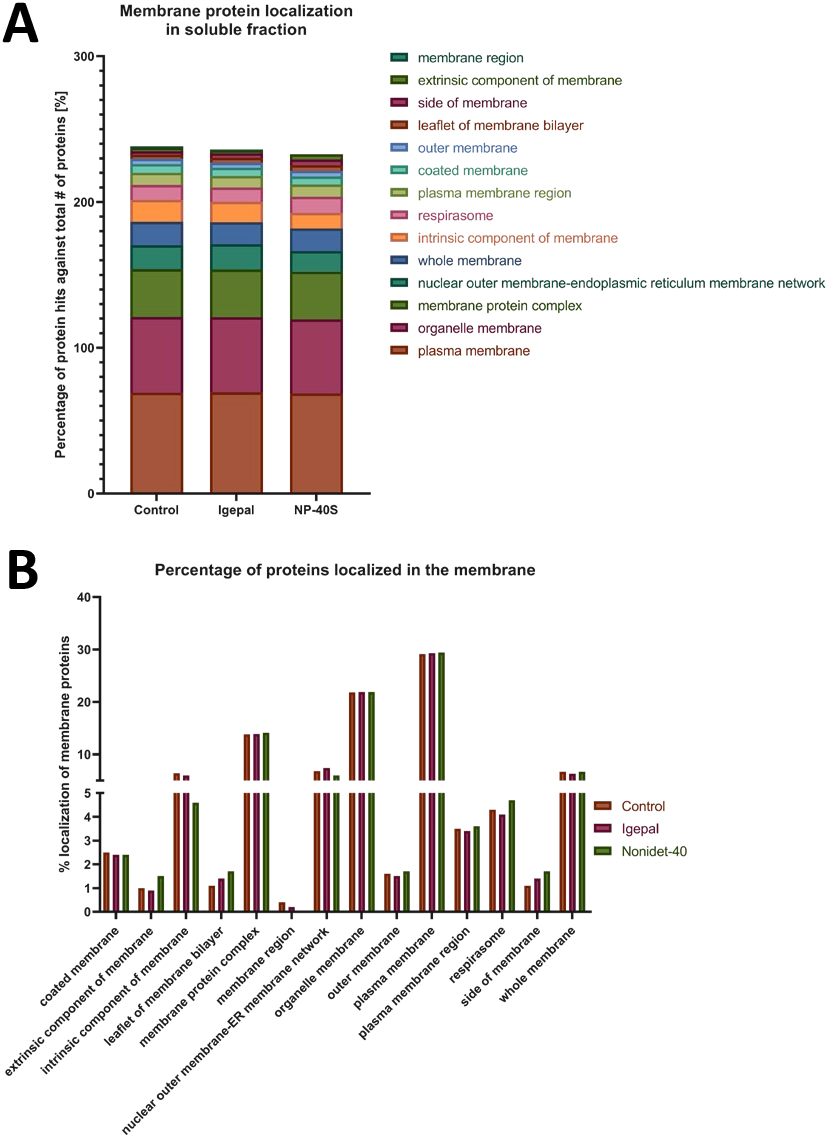
Analysis of distribution of membrane proteins in the soluble fractions after solubilization with different extraction solutions. (A) Bar diagram representing the distribution of membrane protein in the soluble fraction based on G0 categories in and (B) Bar diagram representing changes in the distribution of membrane proteins in different subcellular locations. The extraction solutions were control PBS, 0.4 % (v/v) Igepal in PBS, 0.4 % (v/v) NP-40S in PBS. The fractions were analyzed by quantitative. mass spectrometry (LC-MS/MS). Data was classified via PANTHER^21^, following Gene Ontology classification.

These extraction methods barely increased the diversity of membrane proteins in solution although nonionic surfactants above their critical micellar concentration (CMC) have been introduced in the extraction solution. However, there were some changes in the composition including a few additional integral membrane proteins in solution. The minimal increases of the membrane protein solubilization rate is not completely unexpected. The utilizing similar solubilization solutions for TPP analysis with samples from K562 cell rendered 18 % of membrane proteins in solution^1^. The conditions to facilitate that a membrane protein would embed in a micelle are complex and required convergence many more factors in addition to the surfactant concentration in the extraction solution. It could require trans-bilayer movements that could be relatedly slow^23^. Based on the concentration of surfactant in solution, and the marginal incorporation of membrane proteins, our solubilization results showed a low membrane protein occupancy of surfactant micelles. It implies the presence of a large proportion of empty micelles and monomers if these soluble proteomes would be studied by TPP analysis. It is well studied that surfactants bind to proteins in solution mainly by electrostatic, hydrophobic, and h-bonding^15,24^. It could be expected that several populations of surfactant from monomers, micelles to self-assembly aggregates will be available for possible interaction with proteins^15^.

### TTP analysis and effect of the temperature on the alteration of protein solubility in aqueous micellar solutions

Analyzing TPP proteomes solubilized by the previously described conditions, the occurrence of interactions of globular or membrane protein with different subpopulation of surfactant molecules could not be excluded based previous results. The principle of TPP method for the identification of targets relays on the detection of the alteration of protein stability under thermal stimulus caused by protein-chemical compound interactions^22^. Therefore, we aimed to evaluate if the interaction between surfactant and proteins could also be detected in the TPP methodology as alteration of protein stability in solution^25^. First, we performed the TPP experiment and evaluated the alteration in protein solubility along the thermal shift range of temperatures in the different solubilization conditions. A total of 3340 proteins were identified in the samples and then filtered according to the requirements for TPP analysis^26^. In the control sample in PBS, the number of proteins identified at the lowest temperature (37 °C) were 1436, and 1738 and 1608 in Igepal and NP-40S respectively. Considering that the surfactant-protein interactions will respond to different principles for globular than membrane proteins, the analysis of alteration in solubility has been presented separately (Fig. 2).

**Figure 2.**
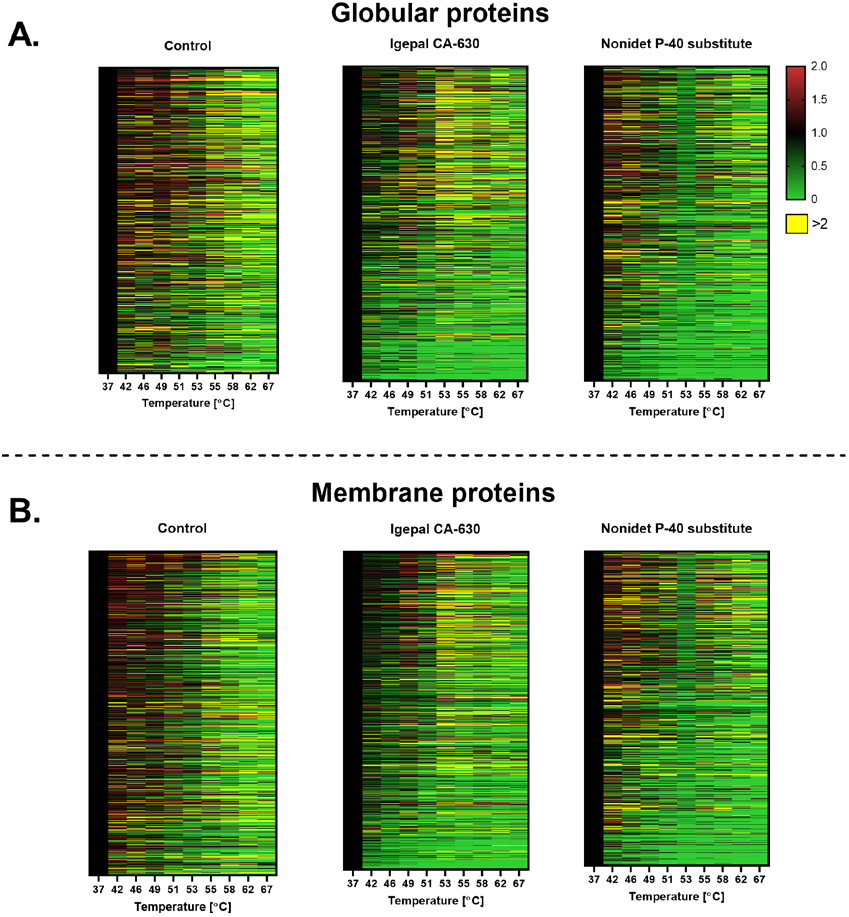
Heatmap representing the effect of the temperature on the alteration of protein solubility in aqueous micellar solutions. The soluble factions were separately exposed to a range of temperature as in TPP methodologies. The fractions were analyzed by quantitative mass spectrometry (LC-MS/MS). The proteins were classified by globular (A) or (B) membrane protein, via Uniprot using Gene Ontology classification. The x-axis represents the temperatures and y-axis represent proteins and each row is one protein. Proteins were normalized against the lowest temperature (37 °C). Each value is the mean of two replicates.

For globular proteins in Igepal solution, proteins are destabilized at the lower temperature than control. It was visible at the heatmap an erratic pattern of proteins changing in solubility from 53 °C to 67 °C. The samples in NP-40S, surfactant with a CPT at 53 °C, a minimal impact is detected at lower temperature but a sharp decrease at CPT, 53 °C. For the membrane proteins in Igepal, the decreased in protein solubility is higher than in control along the thermal shift range. For NP-40S, similarly to the globular proteins, the impact of the phase separation around the CPT temperature marked a decrease in solubility that is higher than at any other temperature of the studied range. In summary, the alteration in protein solubility of proteins in Igepal solution followed erratic changes. In the case of NP-40S, the impact of the CPT was denoted in the sharp effect in protein destabilization at that temperature (Fig. 2).

The results showed that the presence of nonionic surfactant in the extraction solution has an overarching effect on the proteome, altering the protein solubility for both, globular and membrane proteins. This effect is more prominent in the proximity of the surfactant CPT, that is the temperature where a phase separation between high and low surfactant concentration is expected to be formed^27^.

### TPP analysis determines the shift of protein melting point (Tm)

The alteration of the protein solubility observed in previous experiment could suggest variation in the protein Tm. The alteration of the Tm is the factor used in TPP to identify protein targets. Therefore, we performed TPP analysis to detect differences in melting point, plotted in Fig. 3. All comparisons started with 3340 proteins and melting curves were later narrowed down based on four quality criteria as described in the method^26^. For this TPP analysis the control was the equivalent to the sample with vehicle, and the solution with surfactant was defined as the interactor, that is the treatment, according to the TPP R package^22^. The control versus Igepal analysis showed 52 proteins, including 23 membrane proteins that had a difference in Tm (Fig. 3A); in the control versus NP-40S, 68 proteins including 25 membrane proteins has altered their Tm (Fig. 3B). The membrane proteins represent the 44 % and 37% of the proteins with shifted Tm when the aqueous solution contains surfactant micelles. (A list of all proteins, melting points and p-value can be found in the supplementary, Table SI-II). When Igepal was analyzed as the interactor, the globular proteins shifted their Tm to lower temperatures than the control. The membrane proteins had a wider distribution of either increased or decreased Tm. In the case of NP-40S, both globular and membrane proteins showed lower Tm than the control. Changes in composition and turbidity of the protein microenvironment are expected with the increase of temperature and in the proximity to CPT. The primary effect of increasing the temperature is to reduce the degree of structure of water near the micelle surface, facilitating the increases of van der Waals attraction due to the closest contact^28^. This attractive interaction between spheric micelles of certain size facilities a closer contact between micelles and leads to strong spatial changes in the microinvironment^28^. However, it is not easy to model or predict the thermal dependent variation on protein stability in this type of combined proteome that contained globular and membrane proteins. These results showed different pattern in the thermal-dependent alteration of protein solubility for globular, and membrane proteins. Both protein classes in the presence of surfactant are described to follow different principles for their stabilization in solution, and therefore, multiple factors could lead to protein-surfactant interactions^29^.

**Figure 3.**
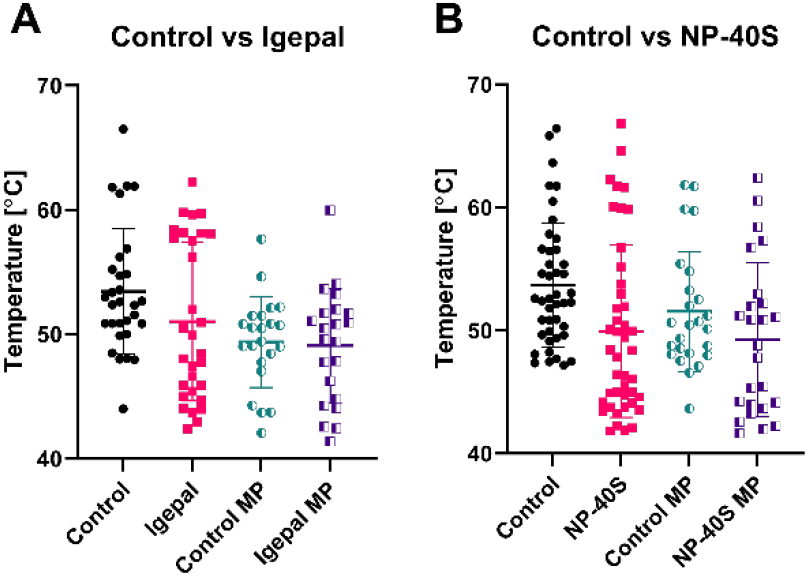
Proteins with shifted melting points (Tm). Proteins with variations in Tm were plotted with a mean melting point. (A) Control set as vehicle for the TPP analysis and Igepal set as interactor. (B) Control set as vehicle and NP-40S set as interactor. In all graphs the middle bar is the mean value of two replicates and the error bars are standard deviation (SD). MP = membrane proteins.

In the case of globular proteins, it should be considered if the responsible for alteration in protein solubility was the interaction with a monomer, a micelle, or other surfactant self-assemblies. In some cases such as human growth factor, individual monomers could bind to hydrophobic patches that could lead to aggregation^30^, and destabilization but on for interferon-γ, the protein is stabilized with the interaction with surfactant^31^. The cooperation of nonionic monomer with micelles has also shown to promote binding to globular proteins and denaturation^32^. Several methodologies could be applied to purified proteins to evaluate the interaction of proteins and small molecules such as limited proteolysis among others^25^. However, it would not be applicable to a proteome-wide approach such as TPP methodologies. These results open the question how the TPP analysis could distinguish between alterations in target solubility based on interaction with surfactant from alteration in solubility due to interactions with a chemical or drug. Therefore, the applicability of the TPP principle of thermal-induced alteration of protein solubility to a proteome extracted with nonionic surfactants could also compromise the robust identification of globular proteins as targets.

In the case of the membrane proteins embedded in micelles, several factors could cause thermal-induced alteration in their solubility. Starting from the solubilization process that included the trans-bilayer movement, a process that is sequential and complex^23^. This process not only required micellization, and the saturation of the membrane bilayer with surfactant, but also the transition of the whole bilayer to thread-like mixed micelles^33^. The thermal-induced changes in the surfactant forms described could also compromise membrane protein solubilization. Secondly, a membrane protein could be embedded in micelles of different sizes. The size of the micelle is a key factor in the precipitation of occupied micelles by sedimentation. The fraction of lipids from the bilayer transferred into the micelles would vary from protein to protein and the same protein could be embedded in micelles of different sizes^23^. The thermal shift would also alter the fluidity of lipids inside the micelle, promoting the micelle size transition^8^. Those micellar structural changes could determine the membrane protein precipitation without even any direct alteration of membrane protein stability. Third, a larger proportion of empty micelles compared to the portion of micelles embedding integral membrane proteins should be expected based on the results. There was a very limited increases in membrane protein solubilization with the nonionic surfactant extraction conditions normally used in TPP^10^. Consequently, empty micelles and occupied micelles should coexist in solution and were available for interactions. Summarizing, it would be very difficult to determine that any observed alteration in solubility of membrane protein embedded in micelles is exclusively caused by a protein-chemical interaction. It should be considered that in case of chemical-protein interaction to membrane proteins, it would take place on a minor hydrophilic domain. This type of domains frequently offers some degree of disorder and flexibility. However, the larger part of membrane protein, the hydrophobic domain is expected to stay protected inside a large surfactant micelle that has shown to be very sensitive to thermal-induced alteration that compromise membrane protein solubility.

### Analysis of aqueous micellar solution of nonionic surfactant used to solubilize membrane proteins for TPP analysis

After evaluating proteomes extracted by nonionic surfactant solutions and determining a thermal dependent alteration of protein solubility, we aimed to analyze the effect of the temperature in the surfactant solutions. The Igepal and NP-40S solutions were prepared at the concentration and procedure utilized for TPP analysis that we have previously described. The DLS measurement from these surfactant solutions was performed at 10 different temperatures between 37 °C to 67 °C, as used for TPP. The images from the DLS wells, and DLS measurements showed thermal dependent changes in size and aggregation of the surfactant in solution.

The DLS well images from the NP-40S solution at 49 °C started to show visual indication of structural changes and turbidity but at lower temperatures variation cannot be visually detected. The instrument could not take images at temperatures higher than 49 °C for safety reasons (Fig. 4A). In addition, a different experiment simulating the heating and subsequently cooling step from the TPP protocol was performed with both, the surfactant extracted proteomes (Fig. 4B), and with surfactant solutions alone (Fig. 4 C, D). The proteomic samples were heated at 55 °C, and 62 °C and cooled to R.T. The surfactant solutions were first heated until 55 °C and 67 °C and cooled at 21°C. Images from the tubes containing the soluble proteomes revealed changes in the turbidity after heating and cooling (Fig 4.B). Similarly, the images from DLS wells from the surfactant solutions still showed turbidity after the cooling (Fig. 4C; D).

**Figure 4.**
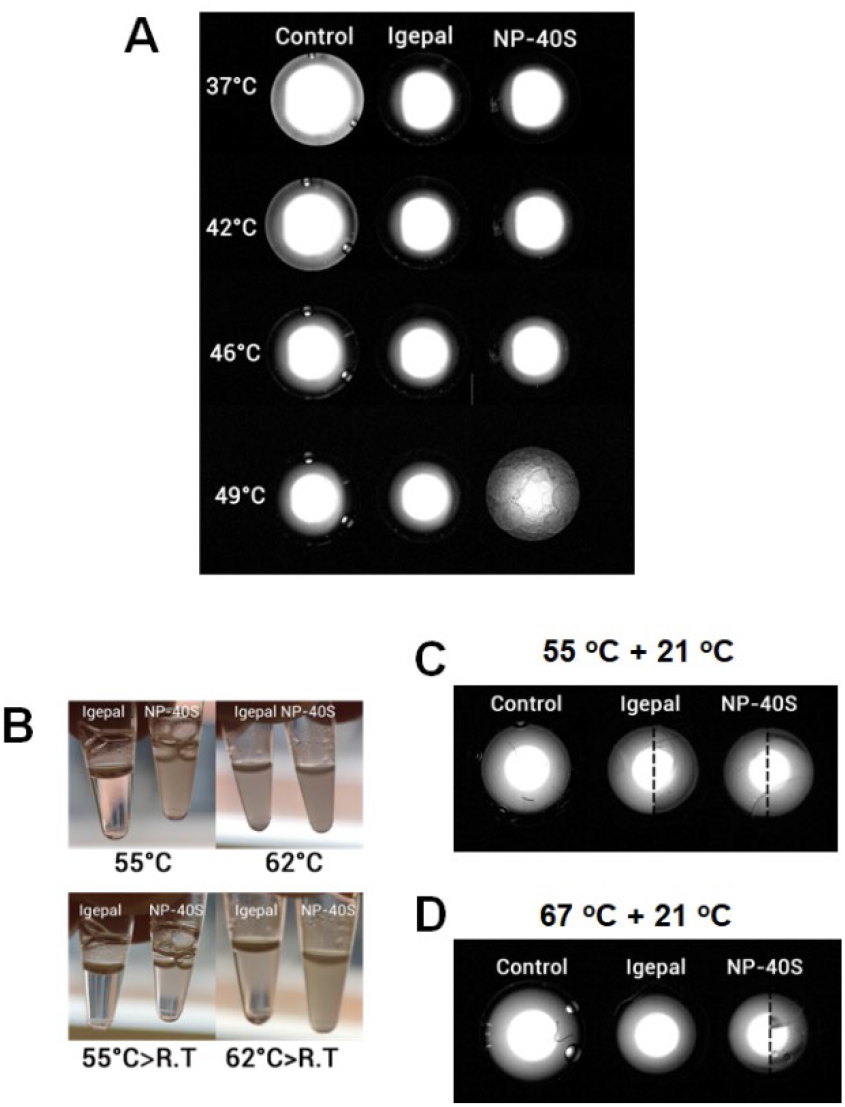
Images from thermal alteration in surfactant solutions, and proteins extracted with surfactant solutions. (A) Images from the bottom of the DLS wells heated at 49 °C. Samples left-to right, control PBS, 0.4 % Igepal in PBS, and 0.4 % NP-40S in PBS. (B) Images from tubes containing the soluble proteome extracted with Igepal (left), and NP-40S (right) after different thermal treatments: heating to 55 °C, heating to 62 °C, heating to 55 °C and cooling to R.T, and heating to 62 °C followed by cool to R.T. (C) Images from DLS wells control PBS versus the 2 surfactant solutions heated to 55 °C and then cooled to 21 °C. (D) Images from DLS wells control PBS versus the 2 surfactant solutions heated to 67 °C and then cooled to 21 °C. The experiments were performed with and without silicone layering to evaluate any possible effects from evaporation. The samples with longitudinal sections are composites where the left side corresponded to the experiment without silicone layering and the right with covering.

Thermal-induced structural alterations of the surfactant solutions were detected from the DLS measurements. In the case of Igepal, DLS measurement from 37 °C to 53 °C could be correlated with the expected micellar size and its gradual increase up to the CPT. Data points were difficult to registered around 53-55 °C, when the solution was likely entering into clouding. The reported CPT for Igepal by the suppliers were 53-67 °C. The DLS measurement for NP-40S also suggested a thermal-dependent increase in the micellar size that were expected to be slightly larger than Igepal. The missing data point between 48-55 °C for NP-40S due to instrumental error connected to turbidity were also close to the expected CPT around 45-50 °C. Both surfactant solutions showed a sharp increase in the DLS measured size from 55-65 °C that were several orders of magnitude above a micellar size but could correspond to complex surfactant self-assembly behavior^34^ (Fig. 5A and B). The DLS measurement from the heating and cooling experiments with surfactant solutions corroborated the presence of large surfactant self-assemblies that did not revert after 3 min at 21 °C (Fig 5A, B blue circles).

**Figure 5.**
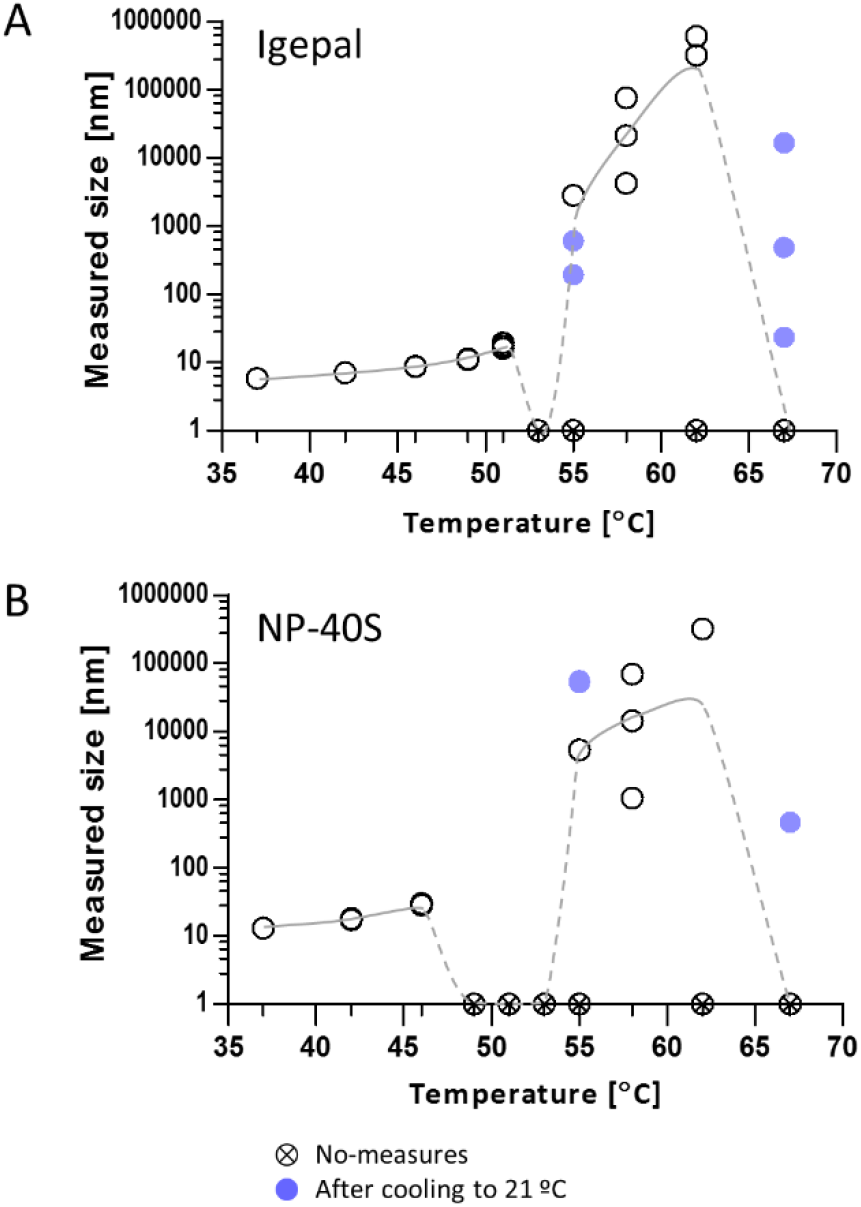
DLS measurement of the nonionic surfactant solutions exposed to 10 stepwise increases of temperature. A: Igepal solutions exposed to 37-67 °C. B: NP-40S solutions exposed to 37-67 °C. White circles represent DLS measurement, crossed circle no-measurement due to auto-attenuating error at the DLS instrument and blue circles were DLS measurement obtained from samples that were separately heated at 55 °C or 67 °C (above CPTs), and cooled to 21 °C. Several data points could not be obtained for technical reasons.

The expected changes on the structure of nonionic surfactant in aqueous solution are defined by the concentration, solution, and the temperature^16^. Our study evaluated the direct effect of the temperature by maintaining constant the rest of the parameters. The surfactant solutions at concentration and the increases of temperature used for TPP analysis showed prominent structural changes. Those changes could be related to micellar growth, clouding, and the accumulation of large structures of surfactant self-assemblies and did not revert after the cooling step.

The solubility of nonionic surfactant in water has been described as a delicate balance between hydrophobic and hydrophilic interactions that assist in maintaining the surfactant molecules soluble. The effects of increase of temperature can easily modify these interactions and decrease the surfactant solubility. Small changes in the effective interaction between surfactant and water cause drastic effects such as phase formation^34^. At clouding the separation between the surfactant rich and surfactant lean starts with changes in turbidity that can be macroscopically observed^16,34^. This is a endothermic transition attributed to conformational rearrangement of the assembled surfactant molecules, specially the polar head groups and its changes in the interaction with water molecules^35^. According to the models proposed, the increase of temperature in the transition phase could lead to a removal of water molecules that were associated with the polar head groups at the bonds, breaking the intermolecular hydrogen bonds. Losing the solvated molecules may induce conformational changes in the micelles with a decrease in volume causing a decrease in stability at the spherical shape. Therefore, the micelles could tend to adopt a more planar arrangement that would stabilize the surfactant molecules in the new environment^16^. The reduction of the micellar curvature could also facilitate association of micelles to micelles with the formation of long cylindrical micelles with bigger volume. Those larger and elongated micelles would require lower centrifugation force to sediment than the smaller micelles that were mainly present at lower temperatures. Therefore, the structural changes of surfactant solutions with the temperature are expected to alter the precipitation pattern of globular proteins interacting with surfactant, and membrane protein in surfactant micelles. It should be reminded that at TPP methods, the centrifugation force is applied to discriminate between soluble and insoluble proteins and to identify the chemical-target interaction.

### Microscropic imaging of proteins exposed to TPP thermal treatment

A microscopic imaging of the proteomes exposed to heat were prepared following the TPP method to visualize microscopical changes in the aqueous micellar solution in the presence of proteins. Samples were heated to 67 °C for 3 min and imaged using light microscopy within the first 3 min, following the heating period required in TPP methodology. In the images, varying sizes of protein aggegates can be observed, as illustrated with the black arrows.

The control contained small protein aggregates located close to each other (Fig. 6A), Igepal sample contained slightly larger protein aggregates (Fig. 6B), and NP-40S contained the largest protein aggregates which are located further apart (Fig. 6C). Those microscopical images showing larger surfactant and protein aggregates confirm that the proteomic samples exposed to the TPP highest temperature followed similar pattern than the surfactant solutions.

**Figure 6.**
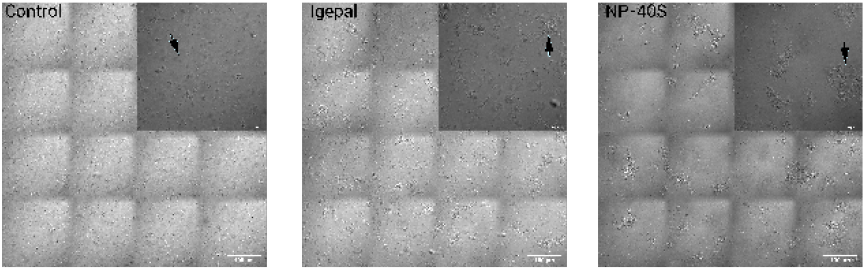
Microscopy visualization of control. Control (PBS) containing proteins following the TPP methodology was heated to 67 °C and visualized within the first 3 minutes. The largest image is merged tiles containing a scale bar of 100 μm. The smaller image is one tile with a scale bar of 30 μm. The black arrow shows one example of a protein aggregate.

## CONCLUSION

The use of nonionic surfactant to increase the solubilization of membrane proteins into the proteome analyzed by TPP for the identification of target of chemicals introduces new parameters that could drastically alter the protein solubility independently from chemical-target interaction. We showed that the standard protocol utilized did not facilitate the solubilization of integral membrane proteins. However, these soluble proteomes in surfactant solutions should contain a large proportion of empty micelles available for interaction with both, globular and membrane proteins, during the TPP protocol. The effects of these interactions were indirectly determined by the broad alteration in protein solubility in the studied proteomes, shift of the protein Tm, and microscopic imaging of proteins forming aggregates. The application of the TPP thermal treatment to the surfactant solution demonstrated that followed the expected structural changes including the increase in the micellar size and formation of other self-assembly structures at the macroscopic size. Our results lead to the exploration of thermal-independent solubilization strategies to assure a confident membrane protein target identification by high-throughput method based in proteome solubility alteration.

## Supporting information

Supplementary material SI and SII. A list of all proteins, melting points and p-value can be found in the supplementary, Table SI-II.

## ASSOCIATED CONTENT

A list of all proteins, melting points and p-value can be found in the supplementary, Table SI-II.

## AUTHOR INFORMATION

### Authors

**Emmanuel Berlin_** Department of Biomedical and Clinical Sciences, Cell Biology, Faculty of Medicine, Linköping University, Linköping, Sweden.

**Veronica Lizano-Fallas_** Department of Biomedical and Clinical Sciences, Cell Biology, Faculty of Medicine, Linköping University, Linköping, Sweden.

**Ana Carrasco del Amor_** Department of Biomedical and Clinical Sciences, Cell Biology, Faculty of Medicine, Linköping University, Linköping, Sweden.

**Olatz Fresnedo-** Department of Physiology, Faculty of Medicine, and Nursing, University of the Basque Country UPV/EHU, Spain

**Susana Cristobal-** Department of Biomedical and Clinical Sciences, Cell Biology, Faculty of Medicine, Linköping University, Linköping, Sweden.

## FUNDING

This work has been performed with funding from: the ERA-NET Marine Biotechnology project CYANOBESITY that it is cofounding from FORMAS, Sweden grant nr. 2016-02004 (SC); the project GOLIATH that has received funding from the European Union’s Horizon 2020 research and innovation programme under grant agreement No 825489 (SC); IKERBASQUE, Basque Foundation for Science (SC); Basque Government Research Grant IT-971-16 and IT-476-22 (SC); Magnus Bergvalls Foundations (SC), VINNOVA No 2021-04909 (SC), and the grant for doctoral studies OAICE-75-2017 World Bank counterpart - University of Costa Rica (VL-F).

## ACKNOWLEDGEMENTS

All the mass spectrometry analysis has been performed with instrumentation at the LiU MS facility.

## CONFLICT OF INTEREST

No.

## LIST OF ABBREVIATIONS

MS: Mass spectrometry
TPP: thermal proteome profilingnonyl-phenyl-polyethylene glycol
NP-40S: (NP-40)octyl-phenoxy(polyoxyethylene)ethanol
Igepal: Igepal CA-630
CPT: cloud point temperature
DLS: dynamic light scattering
Tm: melting point
TSA: thermal shift assay

## Notes

### Competing Interest Statement

The authors have declared no competing interest.

