## Supplementary material SI and SII. A list of all proteins, melting points and p-value can be found in the supplementary, Table SI-II. for "Nonionic surfactants can modify the thermal stability of globular and membrane proteins interfering with the thermal proteome profiling principles to identify protein targets"

### Supplementary data

#### 1. Differences in melting point

**Table SI. Control versus Igepal.** Melting points of control versus Igepal deemed to have significant differences in melting point (°C).

| Protein ID | p-value | Control 1 melting point | Control 2 melting point | Igepal 1 melting point | Igepal 2 melting point | Membrane protein? |
|---|---|---|---|---|---|---|
| A0A0G2JSL8 | 0,00540185 | 50,7767051 | 50,9221086 | 51,9050435 | 51,4575308 | Yes |
| A0A0G2JSU8 | 0,00796226 | 50,4120043 | 48,4553987 | 53,5108695 | 53,7539952 | Yes |
| A0A0G2JVM0 | 0,00127693 | 46,7404893 | 48,7696432 | 42,6333378 | 42,5210238 | Yes |
| A0A0G2K3K2 | 0,0118378 | 48,9456977 | 49,2034033 | 45,0669228 | 44,5334136 | Yes |
| D3ZGY4 | 0,00510369 | 51,7112668 | 51,2800315 | 46,1579154 | 46,3498469 | Yes |
| F1LPV8 | 0,01377531 | 49,8566479 | 48,1839852 | 51,6192578 | 51,793988 | Yes |
| O88618 | 0,01594149 | 50,7567677 | 50,7034299 | 46,7246265 | 48,8134996 | Yes |
| P02680 | 0,01223651 | 52,2673897 | 51,9809523 | 46,0665414 | 49,5868119 | Yes |
| P08011 | 0,00630277 | 49,5921586 | 48,4222403 | 43,2053943 | 44,878113 | Yes |
| P10634 | 0,00264906 | 46,8016468 | 50,1039276 | 41,0376737 | 43,8781869 | Yes |
| P11711 | 0,04747374 | 41,5525948 | 42,603827 | 49,5443566 | 49,2567707 | Yes |
| P16617 | 0,00147144 | 50,8511714 | 50,2336488 | 51,777674 | 52,1904541 | Yes |
| P20070 | 0,00061777 | 43,0264641 | 44,3801745 | 49,8438486 | 51,0850389 | Yes |
| P20673 | 0,02766329 | 52,1997995 | 52,186412 | 50,6856355 | 50,8913414 | Yes |
| P34058 | 0,00127693 | 55,4970973 | 53,7703033 | 50,6390839 | 51,5263726 | Yes |
| P63018 | 0,01849382 | 50,8064551 | 50,159342 | 52,2935786 | 54,1110465 | Yes |
| Q09073 | 0,02583379 | 46,8296728 | 47,2242471 | 42,1852758 | 40,5618801 | Yes |
| Q4KMA8 | 0,01272818 | 57,7811357 | 57,5346658 | 48,2453377 | 50,9787914 | Yes |
| Q5M9H2 | 0,0236471 | 51,1988511 | 51,7989867 | 61,9524002 | 57,9914829 | Yes |
| Q66HF3 | 0,01319786 | 45,1384557 | 42,3200999 | 50,8444268 | 50,986118 | Yes |
| Q9WVK3 | 0,00060873 | 50,7758197 | 50,8359233 | 44,9651945 | 43,571371 | Yes |
| Q9Z2L0 | 2,3769E-05 | 43,5281362 | 45,0325477 | 53,034202 | 55,0755031 | Yes |

| | | | | | | |
|---|---|---|---|---|---|---|
| A0A096MIX1 | 0,00097135 | 50,873583 | 50,8649762 | 43,537191 | 41,2811323 | No |
| A0A0G2JSV0 | 0,01361306 | 52,6452924 | 54,0895422 | 62,0807081 | 57,4538862 | No |
| A0A0G2JTL5 | 0,03307392 | 48,0544908 | 48,939493 | 43,0098813 | 46,4116351 | No |
| A0A0G2JXT3 | 1,369E-05 | 51,9400496 | 52,7318757 | 45,9076529 | 45,9218476 | No |
| A0A0G2QC04 | 0,03695056 | 52,9191255 | 54,2118377 | 50,7886813 | 50,1747662 | No |
| B1WC26 | 0,00622572 | 57,0424206 | 56,6808186 | 57,8918605 | 57,0764613 | No |
| D3Z8I7 | 0,00084152 | 52,3551251 | 52,3929324 | 43,2972847 | 47,5148752 | No |
| D3ZUU8 | 0,00020123 | 55,1626323 | 55,2421971 | 50,9223233 | 51,0083696 | No |
| F1LRV4 | 1,7851E-05 | 50,8262712 | 50,8907792 | 44,3741321 | 43,7223565 | No |
| F1LZ34 | 0,02918255 | 50,8833406 | 52,2246626 | 61,8752089 | 62,6282273 | No |
| F7EV94 | 0,0236471 | 47,3277169 | 48,786561 | 45,1060516 | 44,8929935 | No |
| G3V617 | 0,04581213 | 50,965951 | 50,8312147 | 48,0058086 | 48,8765448 | No |
| M3ZCQ0 | 0,01519224 | 47,2202332 | 48,6701561 | 42,8156469 | 44,6002379 | No |
| P24473 | 0,00263864 | 47,0029415 | 49,0034302 | 57,5865762 | 54,8267688 | No |
| P29266 | 0,02922103 | 61,3294696 | 61,2939244 | 57,9441546 | 58,282011 | No |
| P32232 | 4,3287E-05 | 54,6738232 | 57,7490831 | 57,9564291 | 61,423994 | No |
| P55159 | 0,00095654 | 53,5253929 | 51,7053998 | 58,284411 | 58,0028303 | No |
| P57093 | 0,0043998 | 53,1558947 | 52,1700732 | 47,953255 | 43,7797471 | No |
| P57113 | 0,00167983 | 48,2750335 | 51,7611195 | 44,1084026 | 43,8386699 | No |
| P68255 | 0,0132931 | 62,22893 | 61,3901202 | 58,2317246 | 57,1534834 | No |
| P85834 | 0,01377034 | 50,9576564 | 50,7873878 | 47,5254026 | 48,5493177 | No |
| Q07071 | 0,00018007 | 43,565488 | 44,4454497 | 47,795059 | 47,6411667 | No |
| Q3MID4 | 0,00742899 | 54,5962441 | 54,824389 | 45,1906124 | 48,0644182 | No |
| Q3MIE0 | 0,02142063 | 54,9378376 | 54,7348732 | 50,2458865 | 53,7439423 | No |
| Q4QQW3 | 0,03639081 | 61,9639016 | 61,8411886 | 61,1782283 | 58,0839409 | No |
| Q5RKH2 | 0,0118378 | 49,7895186 | 49,9800032 | 45,039143 | 40,8903412 | No |
| Q5XIC0 | 0,00147144 | 52,7662666 | 53,164995 | 49,5688089 | 50,2340613 | No |
| Q68FR6 | 0,00492822 | 50,8418807 | 51,3354364 | 46,9493324 | 47,9046059 | No |

| | | | | | | |
|---|---|---|---|---|---|---|
| Q8K3R0 | 0,00852123 | 66,65881 | 66,3354331 | 58,1546209 | 58,0121648 | No |
| Q8VHT6 | 0,01594149 | 61,7366091 | 62,0612877 | 58,330955 | 58,4931206 | No |

**Table SII. Control versus NP-40S.** Melting points of control versus NP-40S deemed to have significant differences in melting point (°C).

| Protein ID | p-value | Control 1 melting point | Control 2 melting point | NP-40S 1 melting point | NP-40S 2 melting point | Membrane protein? |
|---|---|---|---|---|---|---|
| A0A0G2JVM0 | 0,017047106 | 46,29792153 | 48,75315057 | 43,7061363 | 43,62778662 | Yes |
| A0A0G2JYB1 | 0,014999614 | 61,88253914 | 61,74504132 | 57,01708047 | 57,58497837 | Yes |
| A0A0G2K3K2 | 0,000513642 | 48,346375 | 49,20348645 | 43,1000912 | 45,3561162 | Yes |
| F1LPV8 | 0,002765123 | 49,32669515 | 48,11897354 | 52,72030425 | 51,17620882 | Yes |
| M0R660 | 0,000131941 | 51,93917095 | 50,49689924 | 42,62753003 | 42,51718079 | Yes |
| O88618 | 1,37837E-06 | 50,42486073 | 50,47910635 | 45,6751449 | 45,03355049 | Yes |
| P06761 | 0,000266371 | 59,65733217 | 59,78561377 | 62,70967309 | 62,09524816 | Yes |
| P08011 | 2,17063E-05 | 48,68083528 | 48,40266226 | 41,80940632 | 42,70591815 | Yes |
| P09034 | 0,011240123 | 47,43069335 | 48,88925816 | 43,48708075 | 44,46605629 | Yes |
| P10634 | 0,004002661 | 46,159203 | 49,92652516 | 40,49685379 | 43,42945219 | Yes |
| P12785 | 0,00272625 | 47,41420796 | 48,517845 | 41,69248944 | 44,80330333 | Yes |
| P18418 | 0,002575991 | 60,1398611 | 59,60529956 | 50,31986341 | 54,15559576 | Yes |
| P34058 | 0,001284183 | 56,80569617 | 54,05085095 | 51,43946921 | 50,3301404 | Yes |
| P50399 | 0,006782931 | 52,73665242 | 53,74512939 | 48,19328427 | 49,36850891 | Yes |
| P63018 | 0,001193019 | 50,13211867 | 50,1272875 | 54,00720281 | 51,93452232 | Yes |
| P82995 | 0,000396041 | 54,53245402 | 55,0713913 | 51,03496723 | 51,16256522 | Yes |
| Q09073 | 0,046007918 | 46,08601808 | 47,00027849 | 41,44754163 | 41,82499287 | Yes |
| Q3MIE4 | 0,011179185 | 48,47318105 | 52,96179989 | 57,96511616 | 58,89736975 | Yes |
| Q5M9H2 | 0,00249893 | 51,75487796 | 52,24251349 | 59,11159002 | 62,00333048 | Yes |
| Q5XI73 | 0,032307044 | 61,56343155 | 61,84889672 | 38,77423195 | 56,81098475 | Yes |
| Q64648 | 0,004413651 | 49,86567027 | 44,32481381 | 55,97854771 | 57,46381011 | Yes |
| Q66HF3 | 0,002814444 | 44,98297405 | 42,30804797 | 50,96519366 | 50,73920364 | Yes |

| | | | | | | |
|---|---|---|---|---|---|---|
| Q66HT1 | 0,023481732 | 52,65249304 | 52,37363187 | 50,95720653 | 51,41739096 | Yes |
| Q9WVJ6 | 0,032589104 | 48,41184789 | 50,2949107 | 42,21491887 | 46,1024215 | Yes |
| Q9WVK3 | 0,000131941 | 50,69712056 | 50,6211189 | 45,06567944 | 45,83330939 | Yes |
| A0A096MIX1 | 0,011134753 | 50,87385117 | 50,80795782 | 46,14351094 | 46,53592694 | No |
| A0A0G2JTL5 | 0,000776464 | 47,52551127 | 48,91582266 | 44,10394354 | 43,34458062 | No |
| A0A0G2JXT3 | 0,000235558 | 51,78987169 | 52,48385782 | 45,19889403 | 46,8892425 | No |
| A0A0G2KAV5 | 9,83332E-05 | 48,31971149 | 50,90430672 | 43,18441284 | 43,92966811 | No |
| B2RYW9 | 0,003757005 | 46,42610859 | 47,93082998 | 49,0292915 | 51,10898763 | No |
| D3Z8I7 | 0,002814444 | 52,58547158 | 52,65117006 | 44,10287772 | 44,23040089 | No |
| D3ZIC2 | 2,49758E-05 | 56,53291542 | 56,37427281 | 47,80896283 | 52,03802346 | No |
| D3ZUU8 | 0,003037674 | 55,40819381 | 55,33624126 | 50,62895571 | 48,27511251 | No |
| F1LZ34 | 0,001001435 | 51,58802116 | 52,02763241 | 59,34276425 | 60,76026413 | No |
| G3V6C2 | 0,038510976 | 61,72907671 | 61,79436921 | 58,05876769 | 61,69698082 | No |
| G3V6C4 | 0,013234719 | 48,95520521 | 50,89706161 | 42,02261946 | 45,51993517 | No |
| G3V7C6 | 0,021009336 | 56,80205239 | 52,36344307 | 42,44211363 | 41,21150521 | No |
| I6L9G6 | 0,013289534 | 54,36050814 | 51,51518003 | 46,16510583 | 46,59772971 | No |
| M0RCU5 | 0,000561315 | 57,53680065 | 58,12046521 | 61,5288063 | 61,87623302 | No |
| M3ZCQ0 | 0,00426238 | 46,35908662 | 48,61504167 | 41,25545052 | 42,49164092 | No |
| P10760 | 0,000777629 | 52,07634592 | 53,11644193 | 50,61511223 | 50,15578767 | No |
| P14173 | 2,17063E-05 | 61,70402461 | 61,75188313 | 55,955988 | 54,34296657 | No |
| P22789 | 0,00023322 | 48,7627734 | 47,36055136 | 42,20355402 | 44,28608869 | No |
| P22791 | 3,24917E-06 | 47,68863582 | 47,69148836 | 41,88230464 | 42,24650814 | No |
| P30713 | 0,017605417 | 58,03797118 | 56,90786188 | 56,4921521 | 49,50153724 | No |
| P41034 | 0,010889791 | 55,97371133 | 54,77310744 | 51,07079868 | 50,56856408 | No |
| P48500 | 1,65954E-06 | 60,62652516 | 60,37039253 | 66,79609975 | 66,83049428 | No |
| P50398 | 0,029768656 | 55,32611524 | 53,91314701 | 38,8487819 | 51,15009439 | No |
| P56571 | 0,029768656 | 57,08379813 | 56,02859736 | 61,72556426 | 61,52610533 | No |
| P57093 | 0,028748994 | 53,15797323 | 51,64072375 | 48,04579588 | 47,61728027 | No |

| | | | | | | |
|---|---|---|---|---|---|---|
| P57113 | 0,002995328 | 47,7411341 | 51,56288585 | 42,66429435 | 44,24960623 | No |
| P62959 | 0,038763948 | 58,08400059 | 59,90079126 | 66,83758175 | 62,33999122 | No |
| P85834 | 0,001196963 | 51,03470751 | 50,74461245 | 46,05755 | 43,82088089 | No |
| Q5RKH2 | 0,000412364 | 49,01642589 | 49,72354926 | 44,66586553 | 44,82252847 | No |
| Q5U2S7 | 0,015421117 | 51,37551048 | 53,19132354 | 48,32079541 | 48,5474676 | No |
| Q5XIC0 | 0,0077387 | 52,70485463 | 53,39498371 | 46,87219255 | 49,86450291 | No |
| Q63150 | 0,031854771 | 65,95519883 | 65,69099594 | 61,90698719 | 62,67279813 | No |
| Q66X93 | 0,030836703 | 47,12837091 | 47,79067469 | 38,79032607 | 45,71132261 | No |
| Q68FR6 | 0,02442857 | 50,62160659 | 51,03799693 | 43,92175722 | 47,10142053 | No |
| Q68FR9 | 0,002698201 | 47,16007703 | 47,53036598 | 48,59950661 | 54,98711487 | No |
| Q68FS4 | 0,000169691 | 62,88370195 | 64,36085909 | 56,49492169 | 56,9664055 | No |
| Q68FU3 | 0,024424729 | 51,90832472 | 48,74312842 | 55,51050508 | 52,03120257 | No |
| Q6IMY6 | 0,017047106 | 52,88976081 | 51,57405066 | 43,75078359 | 45,41746837 | No |
| Q6P6U2 | 0,006628626 | 48,30835578 | 49,95750336 | 42,10987539 | 46,07245855 | No |
| Q7TPB1 | 0,027899554 | 54,71083199 | 54,68304761 | 51,94216677 | 51,91327665 | No |
| Q8K3R0 | 0,033484274 | 66,38877112 | 66,45035119 | 58,05955731 | 61,80047385 | No |
| Q920F5 | 0,001294746 | 57,71563646 | 55,51313463 | 38,9090836 | 51,06798976 | No |
| Q9ER34 | 0,032307044 | 54,02142335 | 54,82277592 | 49,43769876 | 50,39161657 | No |
